# Neuron specific ablation of eIF5A or deoxyhypusine synthase leads to impairment in development and cognitive functions in mice

**DOI:** 10.1101/2021.05.11.443636

**Authors:** Rajesh Kumar Kar, Ashleigh S. Hanner, Matthew F. Starost, Danielle Springer, Teresa L. Mastracci, Raghavendra G. Mirmira, Myung Hee Park

## Abstract

Eukaryotic initiation factor 5A (eIF5A) is an essential factor with a unique amino acid, hypusine, required for its activity. Hypusine is formed exclusively in eIF5A by a post-translational modification involving two enzymes, deoxyhypusine synthase (DHPS) and deoxyhypusine hydroxylase (DOHH). Each of the three genes, *Eif5a, Dhps* or *Dohh* is required for mouse embryonic development. Variants in *EIF5A* or *DHPS* were recently identified as the genetic basis underlying certain rare neurodevelopmental disorders in humans. To investigate the roles of eIF5A and DHPS in brain development, we have generated four conditional knockout mouse strains using the *Emx1*-*Cre* or *Camk2a*-*Cre* strain and examined the effects of temporal- and region-specific deletion of *Eif5a* or *Dhps*. The conditional deletion of *Dhps* or *Eif5a* by *Emx1* promotor driven Cre expression (E.9.5, cortex and hippocampus) led to gross defects in forebrain development, reduced growth and premature death. On the other hand, the conditional deletion of *Dhps* or *Eif5a* by *Camk2a*-promoter driven Cre expression (postnatal, mainly in the CA1 region of hippocampus) did not lead to global developmental defects; rather, these knockout animals exhibited severe impairment in spatial learning, contextual learning and memory, when subjected to the Morris Water Maze test and a contextual learning test. In both models, the *Dhps* knockout mice displayed more severe impairment than their *Eif5a* knockout counterparts. The observed defects in brain, global development or cognitive functions most likely result from translation errors due to a deficiency in active, hypusinated eIF5A. Our study underscores the important roles of eIF5A and DHPS in neurodevelopment.

**Significance:** eIF5A, an essential translation factor, is the only protein that undergoes a unique posttranslational modification, that converts lysine to hypusine by conjugation of the aminobutyl moiety from the polyamine spermidine. Hypusine biosynthesis occurs by two enzymatic steps involving deoxyhypusine synthase (DHPS) and deoxyhypusine hydroxylase (DOHH). Mutations in *EIF5A* or *DHPS* have been associated with rare neurodevelopmental disorders in humans. To understand the mechanisms underlying the pathogenesis of the disease, we generated mutant mice with brain-specific deletions of *Eif5a* or *Dhps*. The *Eif5a* and *Dhps* conditional knockout mice exhibited impairment in brain development, growth and cognitive functions. These animal models may serve as useful tools in the development of therapies against the eIF5A- or DHPS-associated neurodevelopmental disorders.

## Introduction

The eukaryotic initiation factor 5A (eIF5A) is the only cellular protein that is activated by a unique posttranslational modification that forms an unusual amino acid, hypusine [N^ε^-(4-amino-2-hydroxybutyl)lysine] (1). Hypusine is essential for the activity of this factor. It is formed in the eIF5A precursor by two consecutive enzymatic steps (Fig 1) (2). The first enzyme, deoxyhypusine synthase (DHPS) (3), catalyzes the transfer of the aminobutyl moiety from the polyamine spermidine to one specific lysine residue of the eIF5A precursor to form an intermediate, deoxyhypusine [N^ε^-(4-aminobutyl)lysine] residue, which is subsequently hydroxylated by deoxyhypusine hydroxylase (DOHH) (4) to complete the synthesis of hypusine (Fig.1). Homozygous, whole-body deletion of any of these three genes, *Eif5a, Dhps* or *Dohh*, in mice causes early embryonic lethality (5, 6) and postnatal conditional deletion of *Eif5a* or *Dhps* leads to inhibition of organ development in mice (7, 8).

**Fig. 1.**
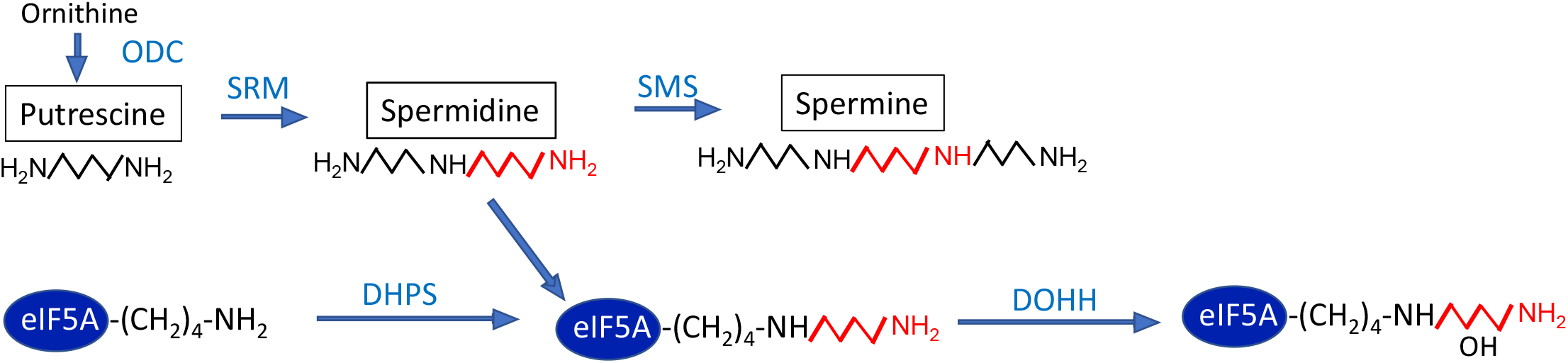
The pathways of polyamine biosynthesis and hypusine modification in eIF5A. The abbreviations are: ODC, ornithine decarboxylase; SRM, spermidine synthase; SMS, spermine synthase; DHPS, deoxyhypusine synthase; DOHH, deoxyhypusine hydroxylase.

Polyamines (putrescine, spermidine and spermine) are essential for eukaryotic cell growth and regulate a vast array of cellular activities (9-11). Polyamine homeostasis is tightly regulated by intricate mechanisms at multiple levels including biosynthesis, catabolism and transport. The majority of cellular polyamines are bound to RNA and the most important function of polyamines appears to be the regulation of translation as polycations (10, 12) and also as a component of hypusine in eIF5A. As hypusine is vital for eIF5A activity and cell proliferation, hypusine synthesis defines a critical function of polyamines in eukaryotic cell growth (13).

In contradiction to its nomenclature, eIF5A facilitates translation elongation rather than translation initiation (14-16). In yeast, eIF5A promotes translation elongation broadly at ribosome stall sites including sequences encoding polyproline stretches and it also enhances translation termination (14, 17). eIF5A binds to the 80S ribosome between the peptidyl-tRNA site and the exit tRNA site (18). The hypusine side chain of eIF5A stabilizes the binding of the peptidyl tRNA to the 80S ribosome and facilitates peptide bond synthesis. Two or more eIF5A isoform genes have been identified in many eukaryotic organisms from fungi to humans. In human and mouse, eIF5A2 shares 84% and 82% amino acid sequence identity with eIF5A1 (usually termed eIF5A). Both isoforms effectively undergo hypusine modification in cells (19). However, only eIF5A1 is constitutively expressed in all mammalian cells and tissues whereas the eIF5A2 isoform mRNA expression appears to be tissue-specific in brain and testis (20). The eIF5A2 protein is normally undetectable in most mammalian tissues and cells, but increased expression of this isoform or eIF5A has been associated with various human cancers (21-23). The *Eif5a2* homozygous knockout mouse develops and grows normally, suggesting that eIF5A2 is dispensible for mouse development (24).

DHPS is known to be totally specific for eIF5A (eIF5A1 and the eIF5A2 isoforms); no other cellular protein is modified by DHPS. The exclusive specificity is based on the requirement for a macromolecular interaction between DHPS and the nearly intact N-domain of eIF5A. A potential role of eIF5A and DHPS in neuronal growth and survival was first suggested in studies that used the neuronal cell line PC12 and rat primary hippocampal cultures *in vitro* (25). In these studies, a reduction of hypusinated eIF5A by using a DHPS inhibitor or DHPS RNAi attenuated neurite outgrowth and neuronal survival (25). Only recently, definitive genetic evidence for their importance in human neurodevelopment was reported (26, 27). From whole genome exome sequencing and genetic analysis, biallelic *DHPS* variants were identified as the cause of a rare autosomal recessive neurodevelopmental disorder (27). More recently, germ line, *de novo*, heterozygous *EIF5A* variants were also reported to be associated with a neurodevelopmental disorder (26). The patients carrying biallelic *DHPS* variants, or heterozygous *EIF5A* variants, share common phenotypes including intellectual disability and developmental delay. Among the five *DHPS* variant patients, four have facial dysmorphism, one has microcephaly and four have clinical seizures. Of the seven *EIF5A* variant patients, all display facial dysmorphism and five of them with microcephaly. Thus, a decrease in the biologically active, hypusinated form of eIF5A appears to interfere with proper neurodevelopment.

To further investigate the roles of eIF5A and DHPS in brain development, we have generated four mouse strains in which either *Eif5a* or *Dhps* is deleted in a temporally and spatially specific manner using the *Emx1-Cre* or the *Camk2a-Cre* line. Phenotype analyses revealed severe morphological defects in the brain, growth retardation and reduced viability in mice with *Emx1-Cre* mediated deletion of *Eif5a* or *Dhps*, and impaired cognitive functions in mice with *Camk2a-Cre* mediated deletion of *Eif5a* or *Dhps*.

## Results

### Generation of four conditional knockout strains: *Eif5a* ^*fl/fl*^*;Emx1-Cre* (*Eif5a*^*Emx*^), *Dhps* ^*fl/fl*^*;Emx1-Cre* (*Dhps*^*Emx*^), *Eif5a*^*fl/fl*^*;Camk2a-Cre* (*Eif5a*^*Camk2a*^) *and Dhps*^*fl/fl*^*;Camk2a-Cre* (*Dhps*^*Camk2a*^)

*Emx1-Cre* mediated knockout of *Eif5a* or *Dhps* was achieved by two-step breeding. First, *Eif5a* ^***fl/fl***^ (8) or *Dhps* ^*fl/fl*^ (7) mouse was mated with *Emx1-IRES-Cre* mouse (28) to generate either *Eif5a* ^*fl/+*^*;Emx1-Cre* or *Dhps* ^*fl/+*^*;Emx1-Cre* mouse, which was mated again with mice carrying their respective homozygous floxed allele to produce either *Eif5a* ^*fl/fl*^; *Emx1-Cre* or *Dhps* ^*fl/fl*^;*Emx1-Cre* mice. *Camk2a-Cre* mediated knockout of *Eif5a* or *Dhps* was achieved as above by the two-step breeding, using the *Camk2a-Cre* transgenic strain T29-1 (29). These four conditional knockout (CKO) mice are referred to as *Eif5a*^*Emx*^, *Dhps*^*Emx*^, *Eif5a*^*Camk2a*^ and *Dhps*^*Camk2a*^, in the rest of the paper. The genotypes of the CKO strains were confirmed by PCR as shown in Fig. S1.

### The effects of temporal- and region-specific knockout of *Eif5a* or *Dhps* in the brain on growth and survival of mice

In the *Emx1-Cre* driven knockout strains, the *Eif5a* or *Dhps* gene is downregulated in the neurons of developing rostral brain including the cerebral cortex, and hippocampus, beginning at E9.5 and continuing throughout postnatal life. On the other hand, in the *Camk2a-Cre* driven CKO strains, the expression of the target gene is abolished postnatally (beginning at P15 – P 21 and continuing through adulthood) in the *Camk2a* expressing neurons in the CA1 regions of hippocampus (29). Differential phenotypes were observed in all four CKO strains. Both male and female groups of *Eif5a*^*Emx*^ pups grew significantly slower than the control *Eif5a*^*fl/fl*^ pups (Fig. 2 *A* and *B*). There was little difference in the growth rates between the male and female groups of the *Eif5a*^*Emx*^ mice, whereas in the control *Eif5a*^*fl/fl*^ group, the males were consistently heavier than the female counterparts (Fig. 2 *A* and *B*). Moreover, survival was reduced in *Eif5a*^*Emx*^ mice compared with control (Fig. 2*C*). The average body weights of both the male and female *Eif5a*^*Emx*^ mice was reduced compared to the control *Eif5a*^*fl/fl*^ mice throughout the period examined (Fig. 2 *A, B, D, E*).

**Fig. 2.**
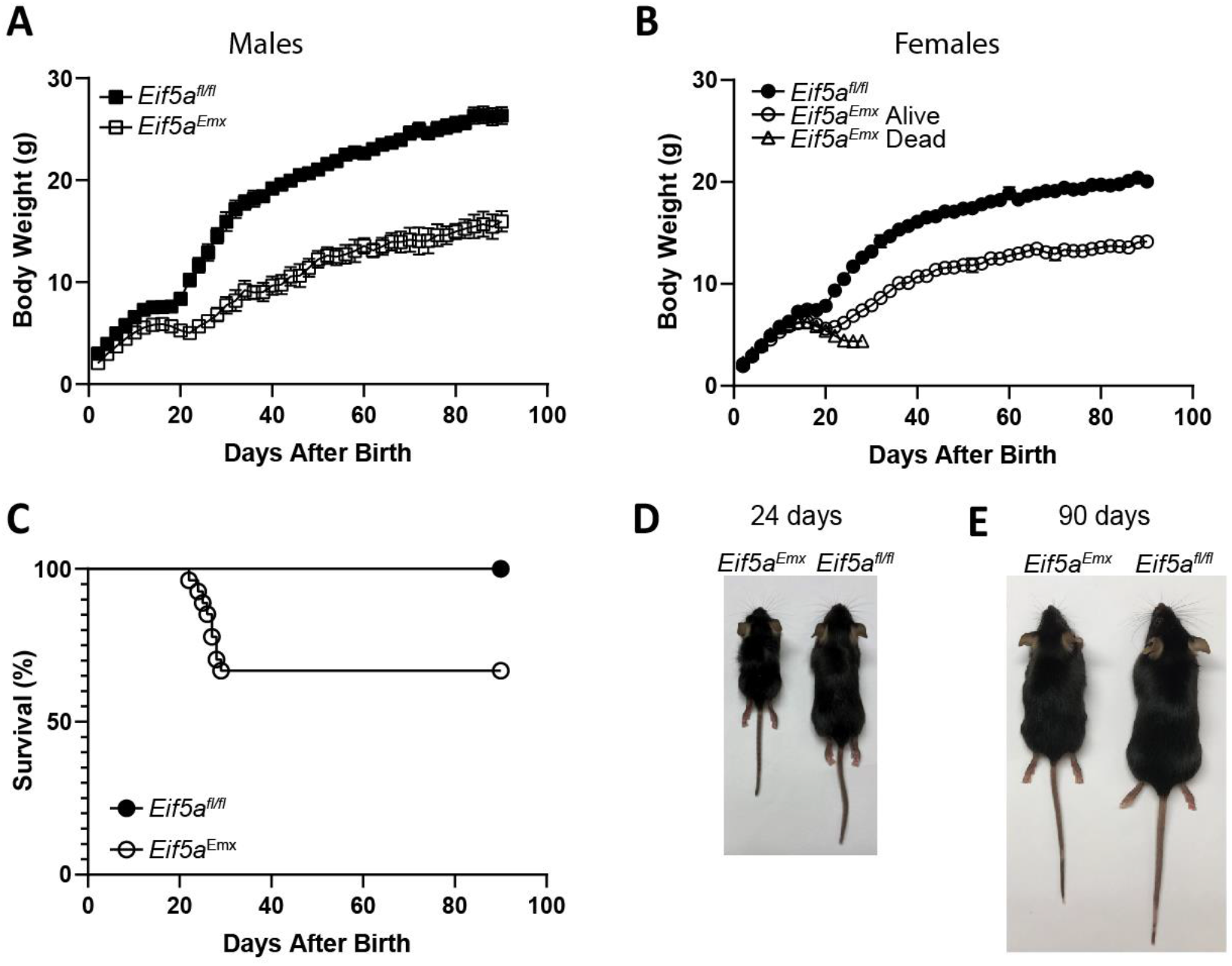
Growth and viability of *Eif5a*^*Emx*^ and the control *Eif5a*^*fl/fl*^ mice. (*A* and *B*) The body weights of the *Eif5a* CKO and the control mice were measured every two days for 90 days and the male and female group body weights are plotted separately in *A* and *B*. The error bars represent SEM. The numbers of mice were *Eif5a*^*fl/fl*^, M (n= 10) and F (n=10); *Eif5a*^*Emx*^, M Alive and Dead combined (n=10), F Alive (n=11) and F Dead (n=6). (*C*) Viability of the *Eif5a*^*Emx*^ and *Eif5a*^*fl/fl*^ mice (male and females combined) in the 90 days after birth. The numbers of mice were *Eif5a*^*fl/fl*^ (n= 20), and *Eif5a*^*Emx*^ (n=27). (*D*) Representative pairs of the *Eif5a* CKO and the control at 24 and 90 days after birth. The body weights of the *Eif5a*^*Emx*^ and the control were 4.03g and 10.99g on day 24 and 13.32g and 21.24g on day 90.

At birth, the *Dhps*^*Emx*^ mice appeared to be similar in size to control mice (*Dhps*^*fl/+*^, *Emx1-cre, Dhps*^*fl/fl*^, *Dhps*^*fl/+*^). However, the postnatal growth of *Dhps*^*Emx*^ mice was significantly impaired (Fig. 3*A*) and nearly arrested by day 12, while the control mice continued to grow. All *Dhps*^*Emx*^ pups died before 4 weeks after birth (Fig. 3*B*). On day 24, *Dhps*^*Emx*^ mice were much smaller than *Dhps*^*fl/fl*^ mice with the average whole body weight less than 50% of the control mice (Fig. 3*C*).

**Fig. 3.**
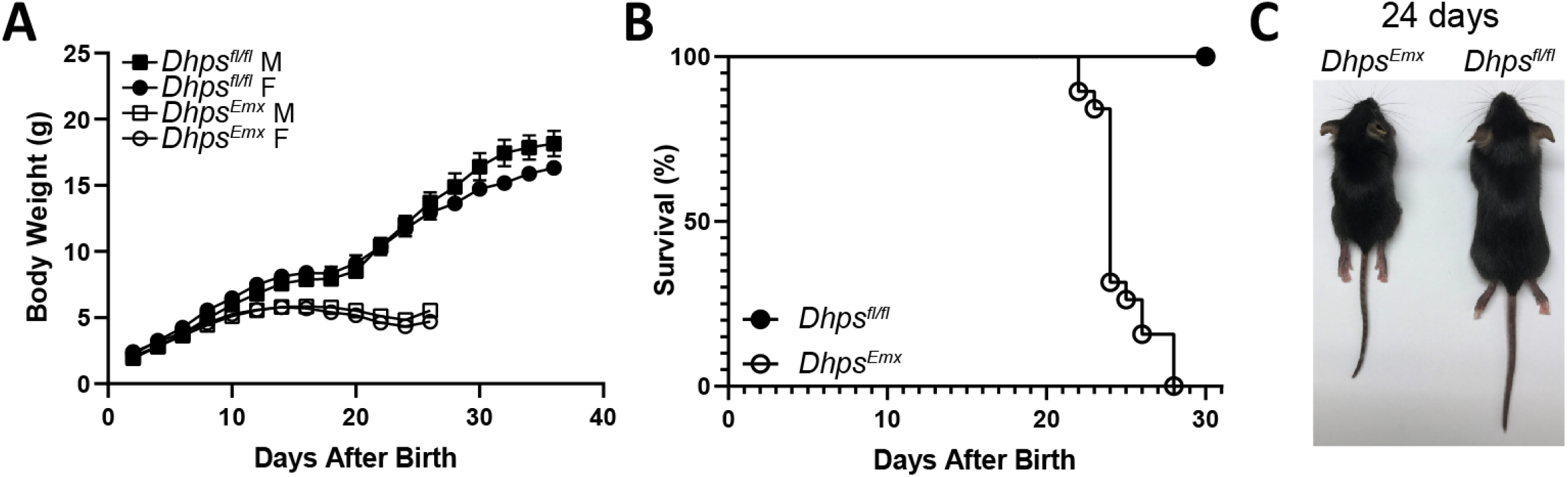
Growth and viability of *Dhps*^*Emxl*^and the control *Dhps*^*fl/fl*^ mice. (*A*) The body weights of *Dhps*^*Emxl*^ and *Dhps*^*fl/fl*^ control, male and female mice were measured every two days starting on day 2 after birth for 36 days until death. The numbers of mice in each group were *Dhps*^*fl/fl*^, M (n= 10) and F (n=10); *Dhps*^*Emxl*^, M (n=11); and F (n=8). The error bars represent SEM. (*B*) Percent survival of *Dhps*^*Emxl*^ and control mice. The numbers of mice in each group were *Dhps*^*fl/fl*^ (n= 20), and *Dhps*^*Emxl*^ (n=19).The n includes both males and females. (*C*) Representative pair of a *Dhps*^*Emxl*^ and a control *Dhps*^*fl/fl*^ mouse on day 24 after birth, with body weights of 5.92g and 10.62g, respectively.

Unlike the deletion of *Eif5a* or *Dhps* in the Emx1 expressing neurons, deletion of either gene in the *Camk2a* expressing neurons did not result in significant inhibition of growth and no visible signs of developmental defects were observed in the first 3 months. However, both *Eif5a*^*Camk2a*^ and *Dhps*^*Camk2a*^ mice lost viability between 2-9 months of age (Fig. S2).

### The effects of deletion of *Dhps* or *Eif5a* on brain development and morphology

The deletion of *Eif5a* or *Dhps* also exerted variable impacts on brain development in the four CKO strains (Figs. 4 and 5). Gross brain images revealed quite similar morphological defects in the brains of *Eif5a*^*Emx*^ and *Dhps*^*Emx*^ mice (Fig. 4 *A* and *C*), even though *Dhps*^*Emx*^ mice displayed more serious defects in growth and survival than *Eif5a*^*Emx*^ mice. The average brain weight of the *Eif5a*^*Emx*^ mice at 4 months was less than that of controls (0.25 g *vs* 0.48 g, respectively). The same gross lesions shown in Fig. 4*A* were observed in all *Eif5a*^*Emx*^ brains (1 and 4 month old mice). The abnormal brain morphology included the loss of the cerebral cortex, hippocampus, corpus collosum, internal capsule and portions of the lateral ventricles and the opening of the third ventricle to the meninges. However, we could not detect cellular changes in the microscopic images of the remaining part of the *Eif5a*^*Emx*^ brain at 4 months (Fig. 4 *I vs J*). The average weight of the *Dhps*^*Emx*^ brains was less than half of the control brain (0.19g *vs* 0.431g) on day 24. Each of the four *Dhps*^*Emx*^ brains examined showed the same gross abnormality (Fig. 4*C*), similar to that of *Eif5a*^*Emx*^ brain (Fig. 4*A*). In the *Dhps*^*Emx*^ brain, the rostral portion of the cerebral cortex was missing or thinned. The deformity also included agenesis of the corpus collosum, hippocampus, internal capsule and the distal portion of the cerebrum overlying the mid-brain. The lumen and the roof of the third ventricle were missing. Microscopic images of remaining *Dhps*^*Emx*^ brain cerebrum showed the neurons enlarged and vesiculated (black arrows, Fig. 4 *K*), not found in the control brain (Fig. 4 *L*).

**Fig. 4.**
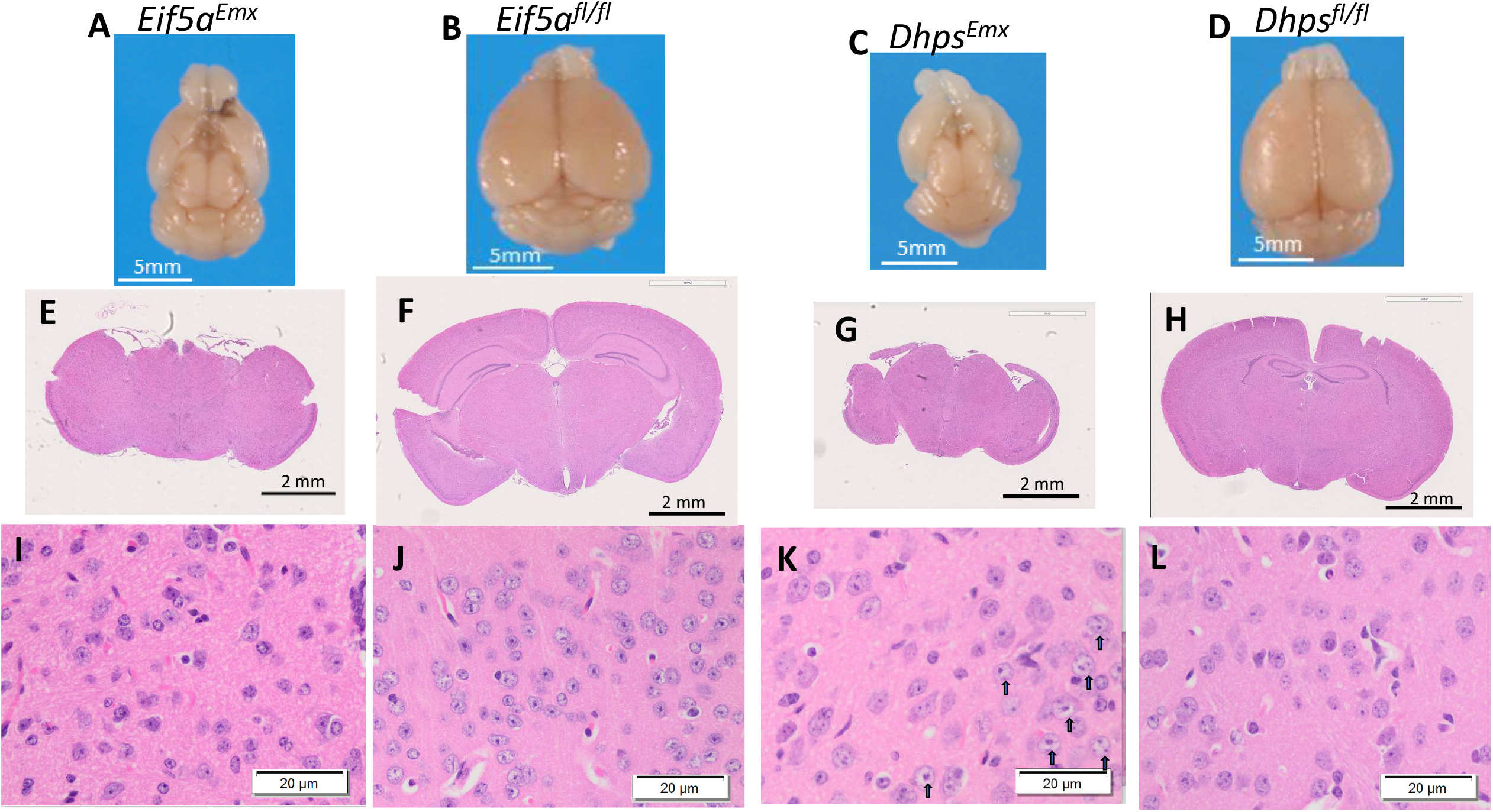
Macro and microscopic changes in the brains of *Eif5a*^*Emx*^ and *Dhps*^*Emxl*^ mice compared to their controls. (*A-D*) A representative whole brain image is shown for each strain: Both *Eif5a*^*Emx*^ and *Dhps*^*Emxl*^ and brains show gross changes in brain size and structures. (*E-H*) A representative coronal section from each brain. (*I-L*) microscopic images of the cortex region of coronal sections show cellular changes including vesiculated nuclei in *Dhps*^*Emxl*^ brains, as indicated by black arrows.

**Fig. 5.**
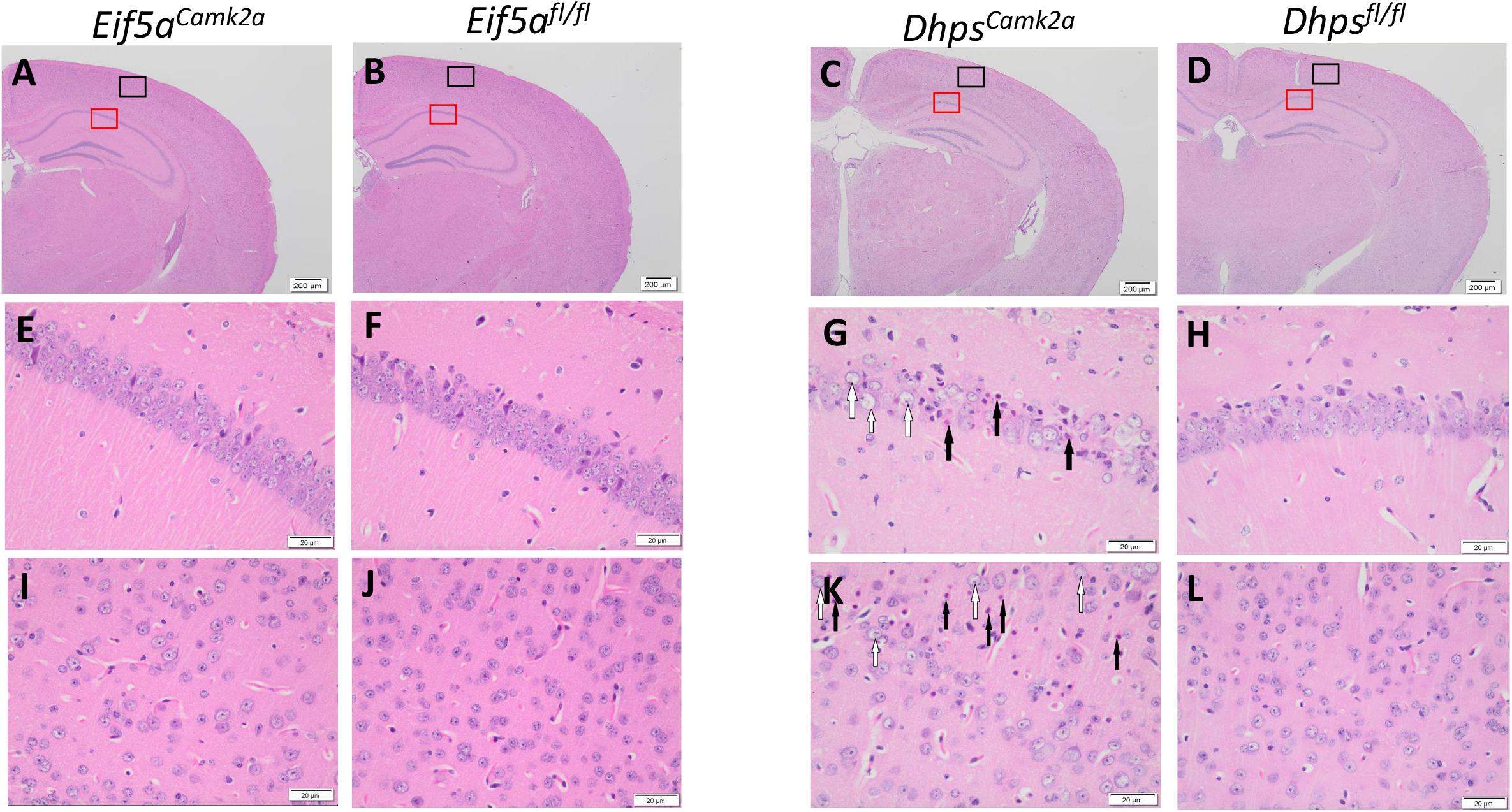
Microscopic images of brain coronal sections of *Eif5a*^*Camk2a*^ and *Dhps*^*Camk2a*^ compared to their controls. (*A-D*) Part of coronal sections of *Eif5a*^*Camk2a*^, *Eif5a* ^*fl/fl*^, *Dhps*^*Camk2a*^, *Dhps* ^*fl/fl*^ mice. (*E-H*) Microscopic images of CA1 region of hippocampus in the red box area. (*I-L*) Microscopic images of external cerebrum in the black box area. Necrotic neurons and vesiculated nuclei are indicated by black and white arrows, respectively.

In contrast to the *Eif5a*^*Emx*^ and *Dhps*^*Emx*^ mice, *Eif5a*^*Camk2a*^ and *Dhps*^*Camk2a*^ mice appeared to grow normally and their gross brain images were indistinguishable from those of control mice (Fig. S3). However, microscopic examination revealed that *Dhps*^*Camk2a*^ mice had extensive neuronal necrosis of the cerebral cortex and hippocampus at 4 months (Fig. 5 *C, G* and *K*). These regions contained necrotic neurons (black arrows) and neurons with enlarged nuclei with extensively vesiculated chromatin (white arrows, Fig 5 *G* and *K*). Similar cellular changes were not observed in the *Eif5a*^*Camk2a*^ brains (Fig. 5 *E* and *I*).

### Impaired cognitive functions in the *Eif5a*^*Camk2a*^ and *Dhps*^*Camk2a*^ mice

We first examined the effects of deletion of *Eif5a* or *Dhps* in *Eif5a*^*Camk2a*^ and *Dhps*^*Camk2a*^ mice on cognitive functions by the Morris water maze test (MWM) (Fig. 6). In this test, the mouse relies on visual cues to navigate to a submerged escape platform. Spatial learning was assessed by daily repeated trials for six days. The latencies to reach the hidden platform for the two controls, *Eif5a*^*fl/fl*^ and *Dhps*^*fl/fl*^, were 37.02 and 38.61 sec, respectively, on day 1 and were shortened to 9.52 and 9.93 sec, respectively, by day 6 (Fig. 6 *A* and *B*). On the other hand, the latencies of *Eif5a*^*Camk2a*^ and *Dhps*^*Camk2a*^ mice on day 1 (42.71 and 53.66 sec, respectively) were longer than those of the respective controls suggesting a poor baseline performance. Furthermore, the improvements of *Eif5a*^*Camk2a*^ and *Dhps*^*Camk2a*^ mice from day 1 to day 6 (reduction of latency by 53% and 38.5%, respectively) were much less than those of the respective control mice (reduction of latency by 74.3% and 75.1%, respectively), suggesting impaired learning in both CKO mice, with *DHPS*^*Camk2a*^ showing greater impairment than *EIF5A*^*Camk2a*^. The analyses of swim distance to the hidden platform also provided a similar indication of learning disability of the two CKO strains (Fig. 6 *C* and *D*). The swim distances were similar for all four groups on day 1, but were significantly longer for the *Eif5a*^*Camk2a*^ and *Dhps*^*Camk2a*^ mice than their respective controls on consecutive days. The improvement indicated by a shortening of the swim distance from day 1 to day 6 was worse for the CKO groups compared to their controls and *Dhps*^*Camk2a*^ consistently underperformed *Eif5a*^*Camk2a*^ mice. These results provide strong evidence that both *Eif5a*^*Camk2a*^ and *Dhps*^*Camk2a*^ mice are impaired in spatial learning and that the impairment is more severe in *Dhps*^*Camk2a*^ than in *Eif5a*^*Camk2a*^ mice (Fig 6. *A, B, C* and *D*).

**Fig. 6.**
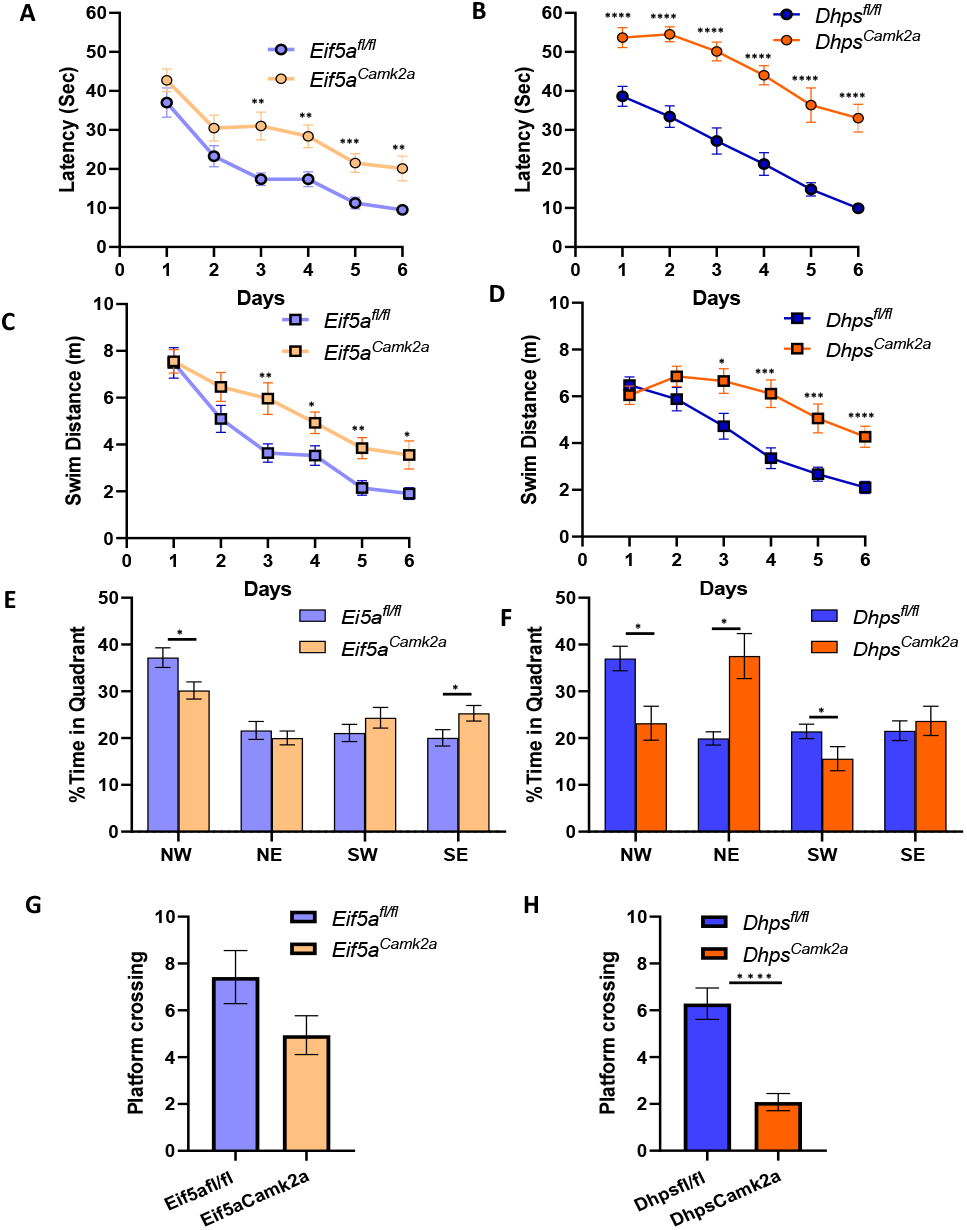
Impaired spatial learning and memory in the *Eif5a*^*Camk2a*^ and *Dhps*^*Camk2a*^ mice. The Morris Water Maze test was performed with the *Eif5a*^*Camk2a*^ and *Eif5a* ^*fl/fl*^ control (*A,C,E,G*) and *Dhps*^*Camk2a*^ and *Dhps* ^*fl/fl*^ control (*B, D, F, H*) as described in the *SI Appendix* Materials and Methods. (*A, B*) The average latency to the platform was significantly longer for the *Eif5a*^*Camk2a*^ (*A*) and *Dhps*^*Camk2a*^(*B*) mice compared to their respective controls. (*C, D*) The average swim distance to the platform by *Eif5a*^*Camk2a*^ (*C*) and *Dhps*^*Camk2a*^ (*D*) mice compared to their controls. (*E, F*) The probe trial measured % time occupancy of *Eif5a*^*Camk2a*^ and *Eif5a* ^*fl/fl*^ (*E*), and *Dhps*^*Camk2a*^ and *Dhps* ^*fl/fl*^ (*F*) in the four quadrants. (*G, H*) The probe trial also measured the number of crossings into the platform area. The numbers of mice in each group were *Eif5a*^*Camk2a*^ (n=17), *Eif5a* ^*fl/fl*^ (n=19), *Dhps*^*Camk2a*^ (n= 13) and *Dhps* ^*fl/fl*^ (n=21),. Error bars represent SEM. \**P* <0.05, \*\**P* <0.01, ***P<0.001, ****P<0.0001 via Student’s t-test.

After the completion of the six day trials, reference memory was evaluated by a probe trial after the removal of the hidden platform. The % time occupancy in the target quadrant (NW) (Fig. 6 *E* and *F*) and and the number of crossings into the small area that previously contained the removed platform were measured (Fig.6 *G* and *H*). The *Eif5a*^*fl/fl*^ and *Dhps*^*fl/fl*^ control groups occupied the target NW quadrant for a significantly longer time, (37.04% and 37.21% time in quadrent, respectively) than in three other quadrants (∼ 20 % in each quadrant). In contrast, the preference to occupy the target quadrant was significantly reduced in *Eif5a*^*Camk2a*^ mice compared to the controls (30.18% *vs* 37.21%) and in *Dhps*^*Camk2a*^ mice compared to the controls (23.20% *vs* 37.04%), suggesting the impaired memory in both CKO groups. The average numbers of crossings into platform area were lower in the CKO mice than in the controls (7.42 and 4.94, respectively, for *Eif5a*^*fl/fl*^ and *Eif5a*^*Camk2a*^ mice and 6.29 and 2.08, respectively, for *Dhps*^*fl/fl*^ and *Dhps*^*Camk2a*^ mice). Both the platform occupancy and the platform area entry data provide clear evidence for memory impairment in the CKO mice, *Dhps*^*Camk2a*^ mice being more deficient than *Eif5a*^*Camk2a*^ mice.

Then a contextual learning (cued fear conditioning) test was carried out as outlined in the top panel of Fig. 7. The baseline freezing and novel context baseline freezing were low and no significant differences were observed among the four groups of mice. However, contextual freezing time was significantly reduced in *Eif5a*^*Camk2a*^ mice (to 48 % of the control *Eif5a*^*fl/fl*^ value) and *Dhps*^*Camk2a*^ (to 40% of *Dhps*^*fl/fl*^ value) (Fig. 7 *A* and *B*). Auditory cued freezing was also reduced in *Eif5a*^*Camk2a*^ mice (to 64% of the control) and in *Dhps*^*Camk2a*^ mice (to 78% of the control), but not as much as the contextual freezing. Taken together, the data in Fig. 6 and Fig. 7 clearly demonstrate the impairment in spatial learning, memory and contextual learning in mice in which *Eif5a* or *Dhps* is deleted in the *Camk2a* expressing neurons of cortex and hippocampus.

**Fig. 7.**
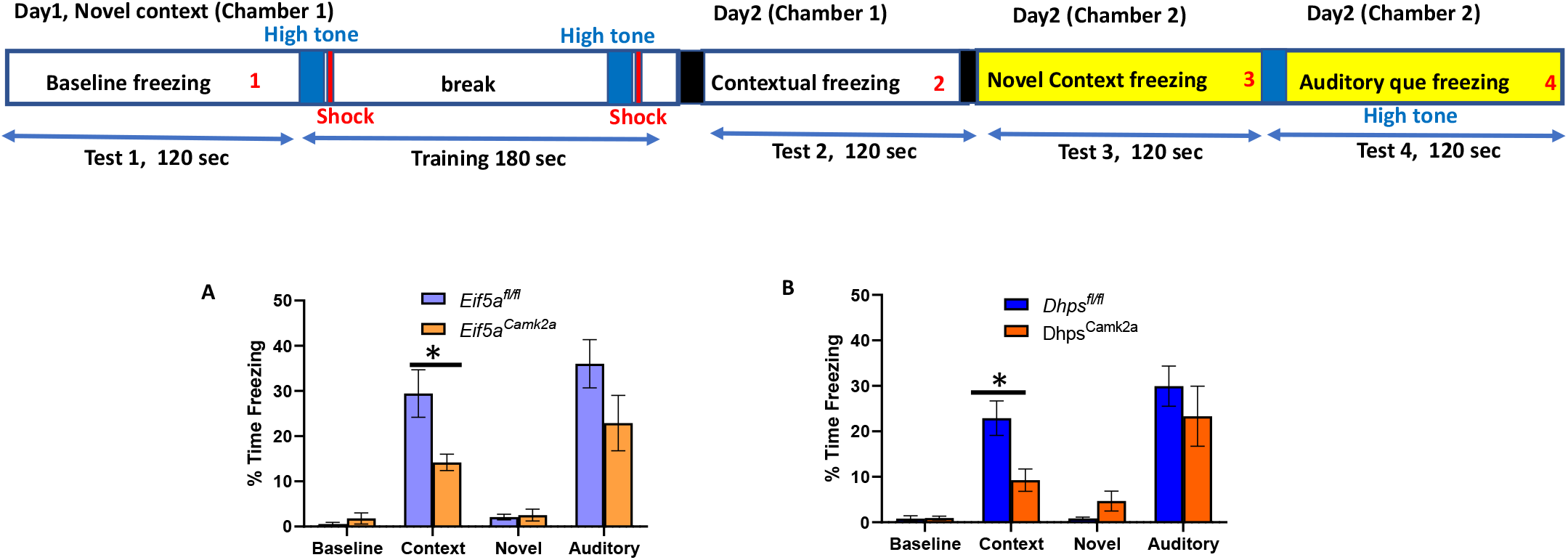
Impaired contextual learning in the *Eif5a*^*Camk2a*^ and *Dhps*^*Camk2a*^ mice. The contextual learning test was performed with the two CKO mice and their control mice, as described in the Materials and Methods. Baseline freezing, contextual freezing, novel context freezing and auditory cue freezing were measured. The numbers of mice in each group were *Eif5a*^*Camk2a*^ (n=11), *Eif5a* ^*fl/fl*^ (n=14), *Dhps*^*Camk2a*^ (n= 9), and *Dhps* ^*fl/fl*^ (n=19).The error bars represent SEM. \**P* <0.05, \*\**P* <0.01, ***P<0.001 via Student’s t-test.

## Discussion

Recent genetic studies have provided evidence that certain variants in the *EIF5A* or *DHPS* gene are associated with rare neurodevelopmental disorders in humans. Furthermore, individuals with *DOHH* variants who display similar developmental delay and intellectual disability have also been identified (Ziegler A. et al, Unpublished results), underscoring the importance of each step of the hypusine modification pathway and thereby the critical role of the hypusinated eIF5A in neurodevelopment in humans. These findings prompted us to generate the four CKO mouse models, with temporal- and region-specific deletion of either *Eif5a* or *Dhps* in the forebrain and to assess the impact on development. Different phenotypes in brain development, growth, survival and cognitive functions were observed in these CKO strains depending on the targeted gene and the Cre-driver. Although the CKO strains do not harbor the same variants of the *EIF5A* and the *DHPS* genes as those of the affected human individuals, it is interesting to note that certain features of the human neurodevelopmental disorders, including intellectual disability, developmental delay, reduced growth, and shortened lifespan, are reflected in the phenotypes of these CKO mice.

The deletion of *Eif5a* or *Dhps* in the *Emx1-*expressing neurons from the mid-embryonic stage resulted in gross morphological abnormalities in brain; cerebral cortex was thinned or missing and hippocampus, corpus collosum and the internal capsule portions of the ventricles were also missing due to agenesis (Fig 4, *A* and *C*). These results indicate that both *Eif5a* and *Dhps* are essential for the embryonic and postnatal development of cortex and hippocampus. Although gross brain defects were similar in the *Eif5a*^*Emx*^ and the *Dhps*^*Emx*^ mice, the deleterious effects on growth and viability were more severe in *Dhps*^*Emx*^ mice than in *Eif5a*^*Emx*^ mice; Approximately 67% of *Eif5a*^*Emx*^ mice survived longer than 3 months whereas all *Dhps*^*Emx*^ mice died within four weeks after birth. The behavioral tests were not performed on these groups of CKO mice because of their short life spans and premature and unpredictable death, especially of *Dhps*^*Emx*^ mice.

The postnatal ablation of *Eif5a* or *Dhps* in the *Camk2a*-expressing neurons did not cause gross changes in the brain compared with those found the *Eif5a*^*Emx*^ and the *Dhps*^*Emx*^ mice, suggesting that the development of cortex and hippocampus was unaltered. Although no significant growth inhibition or visible defects were found in *Eif5a*^*Camk2a*^ or *Dhps*^*Camk2a*^ mice, they both died prematurely between 2-9 months. Furthermore, they displayed concrete evidence of impairment in cognitive functions (Figs. 6 and 7), *Dhps*^*Camk2a*^ being more affected than *Eif5a*^*Camk2a*^ mice. We speculate that this is due to compromised neuronal activities in the hippocampus. Although the genetic alterations in these CKO mice are different from those in the affected human individuals, these cognitively impaired CKO mice hold potential utility in the future development of chemical or biological therapeutics for human neurodevelopmental disorders caused by variants of *EIF5A*, or *DHPS*.

Common phenotypes between *Eif5a*^*Emx*^ and *Dhps*^*Emx*^ mice, and between *Eif5a*^*Camk2a*^ and *Dhps*^*Camk2a*^ mice strongly suggest that a common pathway underlies the impairment in both CKO mice. However, it is hard to explain why the ablation of *Dhps* is more detrimental than that of *Eif5a*, as is evident in all the observed phenotypes. This is counter-intuitive, as the hypusinated eIF5A is the direct player in translation elongation, whereas DHPS is a modifier of eIF5A activity. One possibility may be that eIF5A2 isoform (modified to the hypusine form) can partially compensate for the loss of eIF5A in the targeted neurons of the mouse brain. However, we did not find clear evidence for accumulation of eIF5A2 isoform protein in brain tissues of control or CKO mice (data not shown). Furthermore, commercial eIF5A2 antibodies often crossreact with eIF5A, making it difficult to detect a very low level of eIF5A2 in the presence of abundant eIF5A. The whole-body knockout of eIF5A is embryonic lethal in mouse, suggesting that eIF5A2 is not induced upon knockout of eIF5A and that eIF5A2 cannot compensate for the loss of eIF5A during early embryonic development (6). Another possibility is the interference of eIF5A(Hpu) activities by unhypusinated eIF5A precursors that accumulate upon depletion of spermidine or upon inhibition of DHPS. Although the unhypusinated eIF5A precursors were inactive and did not appear to interfere with the activity of hypusinated eIF5A in the methionyl-puromycin synthesis *in vitro* (30), their potential effects on translation need to be reevaluated *in vivo*. It is possible that the eIF5A precursors still associate with the 80S ribosome through interactions not involving the hypusine residue (18) and interefere with the action of hypusinated eIF5A in mammalian cells and tissues. In such a case, the potential interference by the eIF5A precursors that may have accumulated in *Dhps*^*Emx*^ and *Dhps*^*Camk2a*^ brains could explain their more deleterious phenotypes. In the case of human patients, a heterozygous *de novo EIF5A* variant with partial activities causes clinical phenotypes (26), suggesting that proper neuronal function in humans cannot tolerate even a partial loss (<50%) of eIF5A activity. The detrimental effects of heterozygous *EIF5A* variants may not be simply due to a reduction in active eIF5A, but may also be compounded by the interference by the eIF5A variants. The molecular basis underlying the better survival and performance of the *Eif5a* CKO mice than the *Dhps* CKO mice warrants further investigation.

The implication of variants of *EIF5A* or *DHPS* in human neurodevelopmental disorders is not surprising in view of the fact that variants in a number of other factors in the translation machinery such as alanyl tRNA synthetase (AARS) and eukaryotic translation elongation factors 2 (EF2) and 1a2 (EF1a2) have been associated with neurodevelopmental disorders (31). Errors during translation elongation can lead to production and accumulation of aberrant proteins that are toxic to neural cells. In human individuals with variants in *EIF5A*, or *DHPS*, major clinical symptoms were associated with neurodevelopment (26, 27), suggesting that neuronal systems are most vulnerable to a deficiency of hypusinated eIF5A. Global proteomics analyses provided evidence that depletion of eIF5A in mammalian cells led to endoplasmic reticulum stress, unfolded protein response and upregulation of chaperone expression (32). In addition to these general effects of eIF5A depletion, it is also possible that there are key regulatory factors of brain development that may be specifically dependent on eIF5A for translation. Future studies will be directed towards elucidation of molecular mechanisms underlying these neurodevelopmental disorders stemming from a reduction in bioactive, hypusinated eIF5A and the identification of downstream effectors of eIF5A.

## Materials and Methods

Mouse strains used and detailed methods on mouse maintenance, genotyping, histochemical analysis, Morris water maze test and the contextual learning test are described in *SI Appendix, Materials and Methods*.

## Acknowdgements

We thank Dr. Edith Wolff, Dr. Roman Szabo and Andrew Cho (National Institute of Dental and Craniofacial Research, National Institutes of Health) for helpful advice and suggestions, and Dr. Hans Johansson (Biosearch Technologies, CA) for critical reading of the manuscript, and the National Institute of Dental and Craniofacial Research Veterinary Resource Core for excellent care of mice, and the National Heart, Lung and Blood Institute Murine Phenotyping Core and Morteza Peiravi for conducting the behavioral tests.

This work was supported by the intramural research program of the National Institute of Dental and Craniofacial Research, National Institutes of Health (M.H.P.), grants from the Juvenile Diabetes Research Foundation 5-CDA-2016-194-A-N (T.L.M) and NIH 1R01DK121987-01A1 (T.L.M.), and NIH R01 DK060581, R01 DK125906, and R01 DK105588 (all to R.G.M.).

The authors declare that they have no conflicts of interest with the contents of this article. The content is solely the responsibility of the authors and does not necessarily represent the official views of the National Institutes of Health.

## Author Contributions

M.H.P. designed research, R.K.K, A.S.H, M.F.S. and D.S. performed experiments and/or analyzed the data. T.L.M. and R.G.M. provided the *Eif5a*^fl/fl^ and *Dhps*^fl/fl^ mice and expertise. M.H.P. wrote the paper with input from coauthors.

## References

1. M. H. Park, H. L. Cooper, J. E. Folk, Identification of hypusine, an unusual amino acid, in a protein from human lymphocytes and of spermidine as its biosynthetic precursor. Proceedings of the National Academy of Sciences of the United States of America 78, 2869–2873 (1981).

2. M. H. Park, E. C. Wolff, Hypusine, a polyamine-derived amino acid critical for eukaryotic translation. The Journal of biological chemistry 293, 18710–18718 (2018).

3. Y. A. Joe, E. C. Wolff, M. H. Park, Cloning and expression of human deoxyhypusine synthase cDNA. Structure-function studies with the recombinant enzyme and mutant proteins. The Journal of biological chemistry 270, 22386–22392 (1995).

4. J.-H. Park et al., Molecular cloning, expression, and structural prediction of deoxyhypusine hydroxylase: a HEAT-repeat-containing metalloenzyme. Proceedings of the National Academy of Sciences of the United States of America 103, 51–56 (2006).

5. H. Sievert et al., A novel mouse model for inhibition of DOHH-mediated hypusine modification reveals a crucial function in embryonic development, proliferation and oncogenic transformation. Dis Model Mech 7, 963–976 (2014).

6. K. Nishimura, S. B. Lee, J. H. Park, M. H. Park, Essential role of eIF5A-1 and deoxyhypusine synthase in mouse embryonic development. Amino acids 42, 703–710 (2012).

7. E. M. Levasseur et al., Hypusine biosynthesis in beta cells links polyamine metabolism to facultative cellular proliferation to maintain glucose homeostasis. Sci Signal 12 (2019).

8. L. R. Padgett et al., Deoxyhypusine synthase, an essential enzyme for hypusine biosynthesis, is required for proper exocrine pancreas development. Faseb j 35, e21473 (2021).

9. A. E. Pegg, R. A. Casero, Jr., Current status of the polyamine research field. Methods in molecular biology (Clifton, N.J.) 720, 3–35 (2011).

10. K. Igarashi, K. Kashiwagi, Modulation of protein synthesis by polyamines. IUBMB life 67, 160–169 (2015).

11. A. E. Pegg, Functions of Polyamines in Mammals. The Journal of biological chemistry291, 14904–14912 (2016).

12. S. Mandal, A. Mandal, H. E. Johansson, A. V. Orjalo, M. H. Park, Depletion of cellular polyamines, spermidine and spermine, causes a total arrest in translation and growth in mammalian cells. Proceedings of the National Academy of Sciences of the United States of America 110, 2169–2174 (2013).

13. M. K. Chattopadhyay, M. H. Park, H. Tabor, Hypusine modification for growth is the major function of spermidine in Saccharomyces cerevisiae polyamine auxotrophs grown in limiting spermidine. Proceedings of the National Academy of Sciences of the United States of America 105, 6554–6559 (2008).

14. T. E. Dever, E. Gutierrez, B.-S. Shin, The hypusine-containing translation factor eIF5A. Crit Rev Biochem Mol Biol 49, 413–425 (2014).

15. A. P. Gregio, V. P. Cano, J. S. Avaca, S. R. Valentini, C. F. Zanelli, eIF5A has a function in the elongation step of translation in yeast. Biochemical and biophysical research communications 380, 785–790 (2009).

16. P. Saini, D. E. Eyler, R. Green, T. E. Dever, Hypusine-containing protein eIF5A promotes translation elongation. Nature 459, 118–121 (2009).

17. A. P. Schuller, C. C.-C. Wu, T. E. Dever, A. R. Buskirk, R. Green, eIF5A Functions Globally in Translation Elongation and Termination. Mol Cell 66, 194-205.e195 (2017).

18. C. Schmidt et al., Structure of the hypusinylated eukaryotic translation factor eIF-5A bound to the ribosome. Nucleic Acids Res 44, 1944–1951 (2016).

19. P. M. Clement et al., Identification and characterization of eukaryotic initiation factor 5A-2. European journal of biochemistry 270, 4254–4263 (2003).

20. Z. A. Jenkins, P.G. Hååg, H. E. Johansson, Human eIF5A2 on chromosome 3q25-q27 is a phylogenetically conserved vertebrate variant of eukaryotic translation initiation factor 5A with tissue-specific expression. Genomics 71, 101–109 (2001).

21. X. Y. Guan et al., Isolation of a novel candidate oncogene within a frequently amplified region at 3q26 in ovarian cancer. Cancer research 61, 3806–3809 (2001).

22. M. B. Mathews, J. W. Hershey, The translation factor eIF5A and human cancer. Biochimica et biophysica acta 1849, 836–844 (2015).

23. S. Nakanishi, J. L. Cleveland, Targeting the polyamine-hypusine circuit for the prevention and treatment of cancer. Amino acids 48, 2353–2362 (2016).

24. N. Pällmann et al., Biological Relevance and Therapeutic Potential of the Hypusine Modification System. The Journal of biological chemistry 290, 18343–18360 (2015).

25. Y. Huang, D. S. Higginson, L. Hester, M. H. Park, S. H. Snyder, Neuronal growth and survival mediated by eIF5A, a polyamine-modified translation initiation factor. Proceedings of the National Academy of Sciences of the United States of America 104, 4194–4199 (2007).

26. V. Faundes et al., Impaired eIF5A function causes a Mendelian disorder that is partially rescued in model systems by spermidine. Nature communications 12, 833 (2021).

27. M. Ganapathi et al., Recessive Rare Variants in Deoxyhypusine Synthase, an Enzyme Involved in the Synthesis of Hypusine, Are Associated with a Neurodevelopmental Disorder. American journal of human genetics 104, 287–298 (2019).

28. J. A. Gorski et al., Cortical excitatory neurons and glia, but not GABAergic neurons, are produced in the Emx1-expressing lineage. J Neurosci 22, 6309–6314 (2002).

29. J. Z. Tsien et al., Subregion- and cell type-restricted gene knockout in mouse brain. Cell 87, 1317–1326 (1996).

30. M. H. Park, The essential role of hypusine in eukaryotic translation initiation factor 4D (eIF-4D). Purification of eIF-4D and its precursors and comparison of their activities. The Journal of biological chemistry 264, 18531–18535 (1989).

31. M. Kapur, S. L. Ackerman, mRNA Translation Gone Awry: Translation Fidelity and Neurological Disease. Trends Genet 34, 218–231 (2018).

32. A. Mandal, S. Mandal, M. H. Park, Global quantitative proteomics reveal up-regulation of endoplasmic reticulum stress response proteins upon depletion of eIF5A in HeLa cells. Sci Rep 6, 25795 (2016).

