## Supporting information for "Neuron specific ablation of eIF5A or deoxyhypusine synthase leads to impairment in development and cognitive functions in mice"

From the [1]Molecular and Cellular Biochemistry Section, NIDCR, National Institutes of Health, Bethesda, MD, 20892, USA; [2]Division of Veterinary Resources, Diagnostic and Research Services Branch, National Institutes of Health, Bethesda, MD, 20892, USA; [3]NHLBI Murine Phenotyping Core, National Heart Lung and Blood Institute, National Institutes of Health, Bethesda, MD, 20892, USA; ; [4] Department of Biology, Indiana University-Purdue University-Indianapolis, Indianapolis, IN, 46202; [5]Department of Medicine, The University of Chicago, Chicago, Il, 60637

Running title: Role of eIF5A in mouse neurodevelopment

*To whom correspondence should be addressed: Myung Hee Park: Molecular and Cellular Biochemistry Section, Bldg 30 Rm 3A300, NIDCR, NIH, 30 Convent Drive, Bethesda MD 20892-4340 USA;; Tel: (301) 496-5056.

### Materials and Methods

***Mouse maintenance and sample collection.*** All experimental procedures involving mice were approved by the NIDCR Animal Care Committee and were conducted in accordance with approved protocols. Pups were housed in an animal facility with a 14/10 h light/dark cycle in positive pressure ventilated racks with filtered-top cages (Lab Products Inc., Seaford, Delaware, U.S.A.). Animals were fed autoclavable rodent pellets, NIH-07 Mouse/Rat Diet (Envigo, #7022, Madison, WI, USA) and UV treated ultra-filtered water *ad libitum* throughout the experiments. To help the feeding of small mutant mice, hydrogel and soft foods were offered in petri dishes on the floor of the cages.

***Mouse strains used for brain specific knockout of Eif5a or Dhps.*** The floxed strains, *Eif5a*$^{fl/fl}$ and *Dhps*$^{fl/fl}$ (Dhpstm1.1Mirm/J, stock #034895, Jackson laboratory) were generated as reported (1, 2). Neither the *Eif5a*$^{fl/fl}$ mice nor *Dhps*$^{fl/fl}$ mice showed phenotypic differences compared with their wild type C57BL/6 littermates, and as a result *Eif5a*$^{fl/fl}$ and *Dhps*$^{fl/fl}$ mice were used as controls for the respective CKO strains. The homozygous *Emx1-IRES-Cre* (B6.129S2-Emx1, Jackson laboratory) (3) were viable, fertile, normal in size and do not display any gross physical or behavioral abnormalities. Recombination occurs in approximately 88% of the neurons of the neocortex and hippocampus, and in the glial cells of the pallium starting E.9.5. The Camk2a transgenic strain, T29-1 (B6.Cg-Tg(Camk2a-Cre, Jackson laboratory) used in the study displayed a normal phenotype and the Camk2a-Cre recombinase was expressed in the forebrain, predominantly in the CA1 pyramidal cell layer in the hippocampus postnatally 3-4 weeks after birth (4).

***Genotyping of knockout mice.*** Genomic DNA was isolated from tail biopsies using QIAmp DNA Blood Mini Kit (Qiagen) according to the manufacturer's instruction. For PCR analysis, mouse tail DNA was amplified using JumpStart Taq ReadyMix (Millipore Sigma) using the following programs: For *Dhps* lox PCR, 95°C for 5 min, denaturation at 95°C for 0.5 min, annealing at 58°C for 0.5 min, an extension reaction at 72°C for 1 min for 35 cycles and a final extension reaction at 72°C for 5 min. For *Eif5a* lox PCR, the same program was used as above except the following changes, 94°C for 2 min, 94°C 0.5 min, 55°C 0.5 min. The primers for detection of the *cre* gene (amplified product, 300 bp) were CreF, 5-GGACATGTTCAGGGATCGCCAGGCG-3 and CreR, 5-GCATAACCAGTGAAACAGCATTGCTG-3. The primers for detecting the floxed *Eif5a* locus were *Eif5a*_loxF: 5-CCA CTT GTC CAC GTT TGT CC -3 and *Eif5a*_loxR: 5'-CAA TGC CAA CCA GAT GGA CC -3 (amplified products, lox: 557 bp, WT: 505 bp) and those for the floxed *Dhps* locus were *Dhps* loxF: 5-GTAAACTAGAGTTCTGCGATGGGTGG-3 and *Dhps* loxR: 5-TCAATCTGGTCATAAGGGCACAGG -3 (amplified products, lox: 396 bp, WT: 319 bp).

***Histochemical analysis.*** The animals were euthanized with carbon dioxide. A necropsy was performed, multiple tissues and organs were collected, placed in 10% formalin and fixed for 24-48 hours. The tissues were then processed through a series of alcohols and xylenes and embedded in paraffin. Serial sections (thickness of 5mm) were prepared and stained with 0.1% Hematoxylin and Eosin (H & E).

***Morris Water Maze (MWM) test.*** Learning plasticity and cognitive flexibility were tested by the Morris water maze test (5) with minor modifications. Although the ages of tested mice varied from 2.5-5 months, a mutant and a matching control (with the same or close to the same age) were tested in pairs. The MWM test was carried out in a circular pool (4 ft. diameter, 30 in. high, San Diego Instruments) filled with water which was made opaque with addition of non-toxic white paint (Crayola) and kept at 20-30°C. A small square clear plexiglass escape platform (hidden platform) was placed in the NW quadrant of the tank, 1 cm beneath the water surface. Swim distance, latency to find platform, swim speed, path length to platform, etc., were measured using behavioral tracking software (ANY-maze). The mice received 4 learning trials per day (trials lasted maximum of 60 seconds) on six consecutive days. After learning trials were completed, a probe trial was performed on the last day, in which the platform was removed. The number of crossings over the location in the pool previously occupied by the removed platform and the percentage of time spent in each of the four pool quadrants were measured for 90 seconds.

***Contextual Learning Test***. The test was conducted following guidelines of a published protocol (6) with modifications. While the animal was in the chamber and provided cues, the following data including total freezing time, number of freezing episodes, duration of freezing episodes and latency between stimuli and freezing was collected. Mice were placed in a sound attenuating chamber, 17cm x 17cm x 25cm (w, d, h) with a light and speaker. After baseline freezing was measured for 120 seconds, auditory tones (2 x 4 kHz 80 dB tone) were delivered into the chamber for 15 seconds, followed by a 2 second foot shock (0.85 mA) through the grid floor. After a break for 120 seconds, the tone-shock procedure was repeated, and the mice were returned to their cages. On day 2 ($24^{th}$ hour), the mice were re-exposed to the chamber used on day 1 and contextual freezing was recorded. Afterwards ($25^{th}$ hour), the mice were placed in a completely new chamber and novel context freezing was measured for 120 seconds. Then, auditory cues were applied, and auditory cue freezing was measured.

***Statistics.*** All data are presented as the mean ± standard error mean (SEM) and were analyzed using the software GraphPad Prism 5.0 (GraphPad Software). Multiple unpaired t-test was used for statistical significance analysis. Statistical significance was defined at $P < 0.05$ and presented as *P < 0.05, **P < 0.01, ***P <0.001 ****P < 0.0001.

**Figure legends**

**Fig. S1**. Confirmation of genotypes of *Eif5a* or *Dhps* CKO mice by PCR. The genomic DNA was isolated from ear punch tissues and PCR was performed as described in Materials and Methods using the primer sets designed to identify the *Eif5a* floxed allele (top panel), the *Dhps* floxed allele (middle panel) and *Cre* transgene (bottom panel). The expected sizes of the products are Eif5a Lox, 557 bp; Eif5a WT, 507 bp; Dhps Lox, 396 bp; Dhps WT, 319 bp; Cre, 300 bp, as indicated by arrows.

**Fig. S2.** Reduced survival of *Eif5a*$^{Camk2a}$ and *Dhps*$^{Camk2a}$ mice. (*A*, *B*) The survival curves of *Eif5a*$^{Camk2a}$ (*A*) and *Dhps*$^{Camk2a}$ (*B*) mice show reduced viability of both CKO mice compared to the respective controls, *Eif5a*$^{fl/fl}$, and *Dhps*$^{fl/fl}$.

**Fig. S3.** Macroscopic images of brains of *Eif5a*$^{Camk2a}$ and *Dhps*$^{Camk2a}$ mice in comparison with the respective controls. The brains were taken from *Eif5a*$^{Camk2a}$ and *Eif5a*$^{fl/fl}$ mice (both 8 months old) and from *Dhps*$^{Camk2a}$ and *Dhps*$^{fl/fl}$ mice (both 3 months old).

### References


1. E. M. Levasseur *et al.*, Hypusine biosynthesis in beta cells links polyamine metabolism to facultative cellular proliferation to maintain glucose homeostasis. *Sci Signal* **12** (2019).
2. L. R. Padgett *et al.*, Deoxyhypusine synthase, an essential enzyme for hypusine biosynthesis, is required for proper exocrine pancreas development. *Faseb j* **35**, e21473 (2021).
3. J. A. Gorski *et al.*, Cortical excitatory neurons and glia, but not GABAergic neurons, are produced in the Emx1-expressing lineage. *J Neurosci* **22**, 6309-6314 (2002).
4. J. Z. Tsien *et al.*, Subregion- and cell type-restricted gene knockout in mouse brain. *Cell* **87**, 1317-1326 (1996).
5. C. V. Vorhees, M. T. Williams, Morris water maze: procedures for assessing spatial and related forms of learning and memory. *Nat Protoc* **1**, 848-858 (2006).
6. H. Shoji, K. Takao, S. Hattori, T. Miyakawa, Contextual and cued fear conditioning test using a video analyzing system in mice. *J Vis Exp* 10.3791/50871 (2014).

Fig. S1.

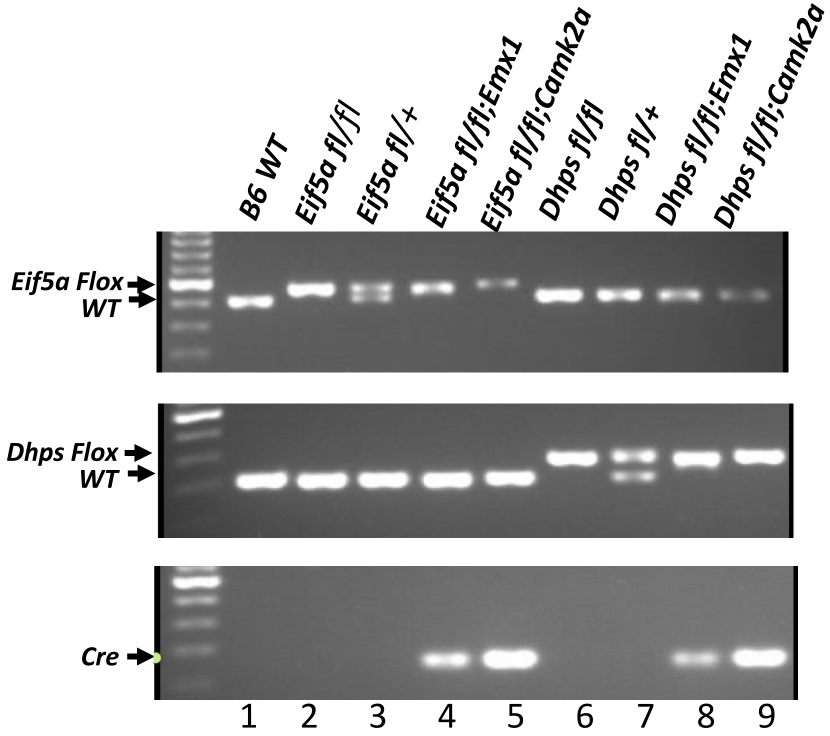

B6 WT
Eif5a fl/fl
Eif5a fl/+
Eif5a fl/fl;Emx1
Eif5a fl/fl;Camk2a
Dhps fl/fl
Dhps fl/+
Dhps fl/fl;Emx1
Dhps fl/fl;Camk2a
Eif5a Flox
WT
Dhps Flox
WT
Cre
1 2 3 4 5 6 7 8 9

Fig. S2.

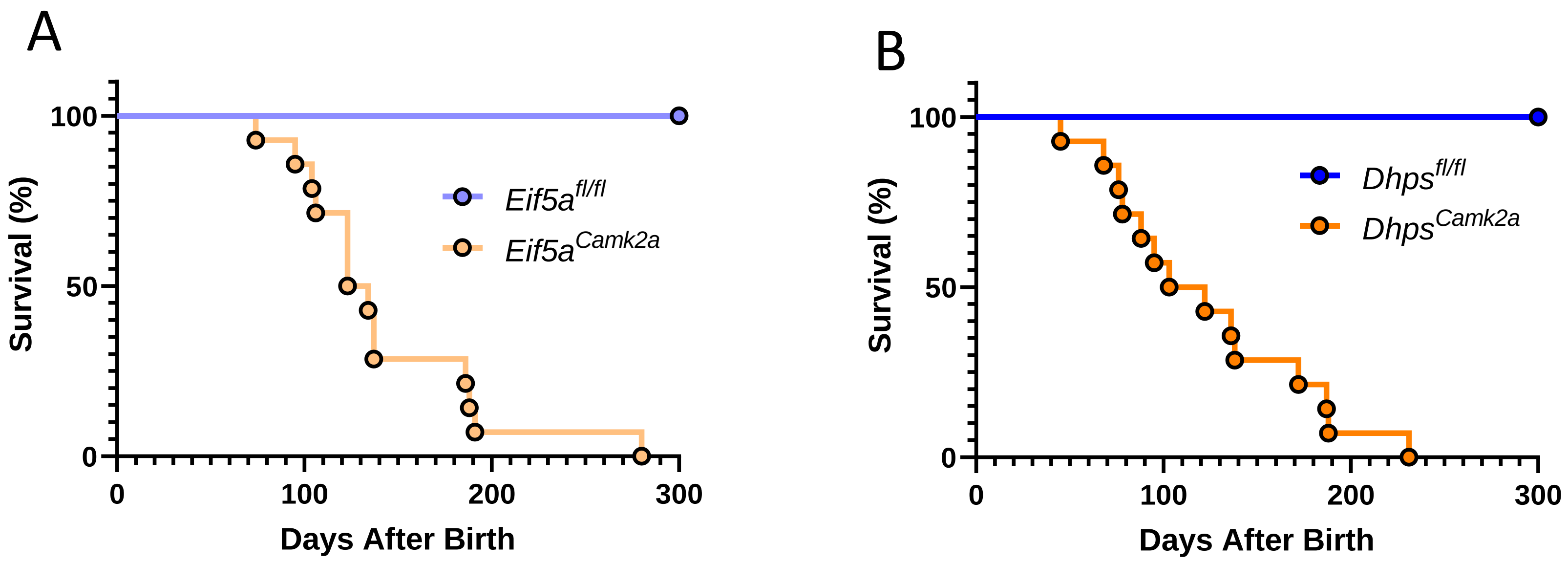

A
Survival (%)
100
50
0
0
100
200
300
Days After Birth
Eif5a^fl/fl
Eif5a^Camk2a
B
Survival (%)
100
50
0
0
100
200
300
Days After Birth
Dhps^fl/fl
Dhps^Camk2a

Fig. S3.

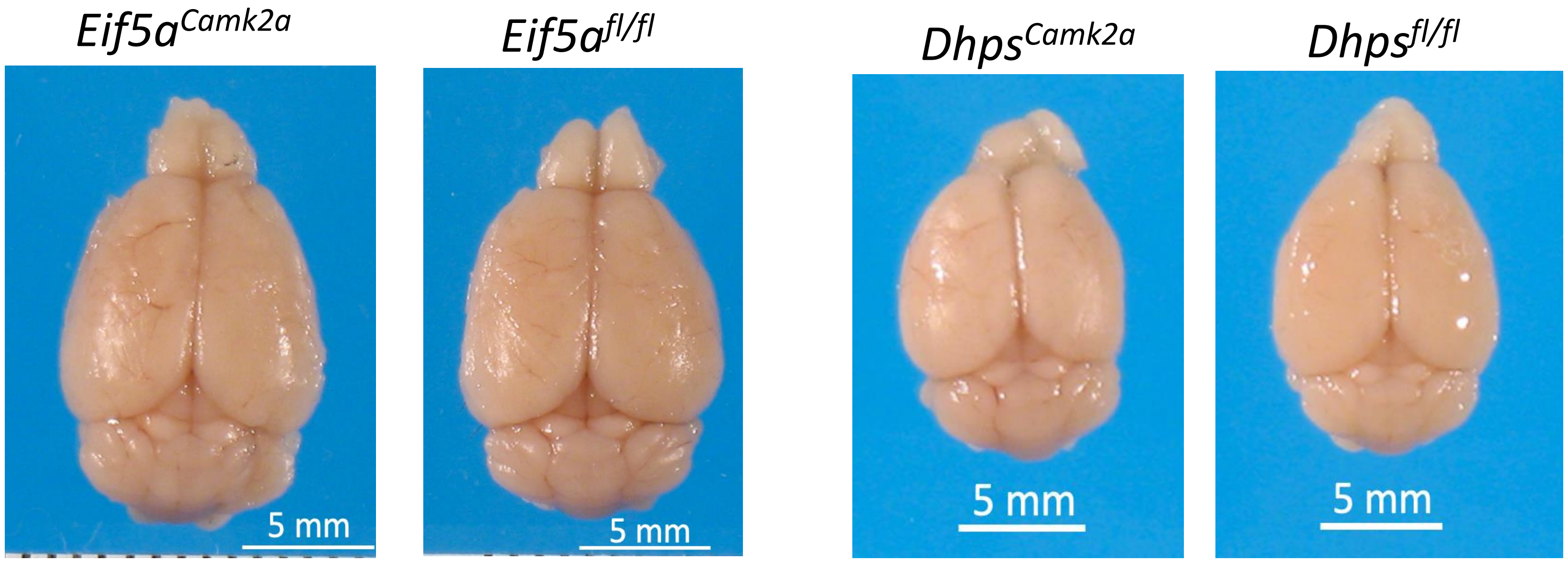

*Eif5a*$^{Camk2a}$
*Eif5a*$^{fl/fl}$
*Dhps*$^{Camk2a}$
*Dhps*$^{fl/fl}$
5 mm
5 mm
5 mm
5 mm
